# PEG-Arginase 1: A Novel Therapy for Optic Nerve Injury

**DOI:** 10.64898/2026.08.26.746813

**Authors:** Mai Yamamoto, Syed A H. Zaidi, Tahira Lemtalsi, Zhimin Xu, Porsche V. Sandow, Robert W. Caldwell, Ruth B. Caldwell, Modesto A. Rojas

## Abstract

Traumatic optic neuropathy (TON) occurs due to direct or indirect injury to the optic nerve and is a significant cause of visual disability. So far, there is no effective treatment. The lack of understanding of the cellular mechanisms by which trauma induces inflammation and damage in retinal neurons is a critical knowledge gap in developing effective therapies. We have studied the role of the arginase 1 (A1) enzyme in this pathology. We have found previously that treatment with a long-acting form of human recombinant A1, pegylated A1 (PEG-A1) after optic nerve crush limits activation of retinal microglia and macrophages (MΦ) and reduces inflammation, thereby decreasing injury and protecting visual function. Here we report on studies designed to demonstrate the therapeutic efficacy of PEG-A1 in mouse models of direct and indirect TON and to elucidate the underlying mechanisms. We used ONC to model direct TON and sonication-induced trauma to the supraorbital rim to model indirect TON (SI-TON). At different times after injury, mice were treated with PEG-A1 which was delivered systemically by i.p. injection or locally by intravitreal injection. In order to assess the role of A1-induced activation of the ornithine/polyamine pathway in the protective effects of PEG-A1, some mice were treated with the ornithine decarboxylase (ODC) inhibitor, difluoromethylornithine (DFMO) immediately after the PEG-A1 treatment. Retinal function was determined by OptoMotry and electroretinography. Retinal injury and microglia/MΦ activation were assessed by immunofluorescence imaging. Expression of inflammatory cytokines was determined by Western blotting and quantitative RT PCR. Liquid chromatography mass spectrometry was used to analyze changes in arginase/ODC pathway metabolites. Results showed that PEG-A1 treatment improved neuronal survival and visual function whether delivered systemically or intravitreally. This neuroprotection was associated with decreased microglia/MΦ activation, decreased inflammatory cytokine expression, and increased formation of L-ornithine and putrescine. Furthermore, DFMO treatment blocked these effects, indicating that PEG-A1 limits retinal injury and preserves vision after ocular injury by activating ODC. ODC processes the arginase product L-ornithine to form polyamines which are known to promote reparative functions. Thus, PEG-A1 therapy offers a new strategy to limit trauma-induced vision loss and promote repair after TON.

## Introduction

Traumatic optic neuropathy (TON) can occur directly due to ocular injury or trauma to the optic nerve, or indirectly due to traumatic brain injury after automobile or bike accidents, falls, assault, or combat injury[1–3] Such injury is often associated with retinal ganglion cell (RGC) death due to primary damage of their axons or injury-related glaucoma [4, 5]. Additional RGC loss can occur secondary to oxidative stress, inflammation, vascular dysfunction, and/or ischemia [5, 6]. So far, there is no effective treatment to limit injury or promote repair. The lack of understanding of the mechanisms of this progressive damage to retinal neurons represents a critical knowledge gap in developing effective therapies. Our recent studies of pathological processes in the eye have focused on role of the urea cycle enzyme, arginase in retinal injury.

Arginase has two isoforms, arginase 1 (A1) and arginase 2 (A2). A2 plays a role in retinal injury during ischemic retinopathy or optic nerve injury, whereas A1 can limit injury in some contexts [7–11]. Our studies of retinal ischemia/reperfusion (I/R) injury showed that A1 deletion globally or in myeloid-derived cells worsens retinal injury and promotes increases in inflammatory microglia or macrophages (MΦ) while suppressing the reparative phenotype [9]. Further studies used treatment with human recombinant A1 linked to polyethylene glycol (PEG-A1) which prolongs its half-life from hours to 3-4 days. PEG-A1 treatment has been used to normalize blood L-arginine levels in patients with A1 deficiency[12]. Pharmaceutical forms of PEG-A1 are currently under development as therapy for cancers that require L-arginine for growth [13–18]. Our *in vivo* studies have shown that PEG-A1 treatment protects against neurovascular injury in models of retinal I/R injury, optic nerve crush (ONC), and stroke by a mechanism involving increases in the reparative microglia/MΦ phenotype and decreases in inflammation[9, 10, 19, 20]. Our *in vitro* studies have shown that PEG-A1 treatment of bone marrow-derived MΦ limits endotoxin-induced inflammation and promotes a reparative phenotype whereas MΦ with A1 deletion show an inflammatory phenotype [9].

Here we have extended our studies to a model of indirect TON and to examine the mechanisms underlying the PEG-A1-mediated neuroprotection after ONC. Arginase hydrolyzes L-arginine to form L-ornithine and urea [21–23]. L-ornithine is metabolized by ornithine decarboxylase 1 (ODC) to form the polyamine putrescine which is further metabolized to form spermidine and spermine [24, 25]. Activation of this pathway can limit oxidative stress and inflammation by several mechanisms. First, arginase-mediated activation of ODC increases polyamine formation which can limit inflammation and promote a reparative MΦ phenotype by increasing putrescine production [26]. Dietary supplement with spermidine has also been shown to limit neuronal inflammatory injury in models of traumatic brain injury, ONC, and glaucoma [27–29]. Additionally, A1 competes with nitric oxide synthase (NOS) for their common substrate L-arginine. During inflammation, high levels of NO produced by upregulation of inducible NOS (iNOS) promote neuronal injury via nitrative/oxidative stress [9]. Thus, increases in A1 can limit injury by reducing iNOS activity [30].

We hypothesized that PEG-A1 limits inflammatory injury during optic neuropathy by activating the L-ornithine/polyamine pathway. We tested this hypothesis by determining the effects of the ODC inhibitor α-difluoromethyl ornithine (DFMO) on the PEG-A1-mediated neuroprotection after ONC. We extended our studies to examine the therapeutic potential of PEG-A1 in limiting retinal injury after indirect TON.

## Materials and methods

### Ethics approval

All animal procedures complied with the Public Health Service Policy on Humane Care and Use of Laboratory Animals (revised July 2017) and with the statement from the Association for Research in Vision and Ophthalmology (ARVO) for the use of animals in ophthalmic and vision research. Procedures were approved by the institutional animal care and use committee (Animal Welfare Assurance # D16-00197). All surgeries were performed under anesthesia and analgesia was provided to minimize suffering. Wild-type (WT) C57BL6J mice (9 – 10 weeks old) were subjected to ONC or TON and sacrificed at various times after the injury.

### Mouse models

Direct traumatic optic neuropathy (TON) was induced by ONC following our previously described method [10]. Indirect TON was induced by using a Branson Sonifier 450 with a 3 mm microtip probe in an acoustic soundproof enclosure chamber as described [31]. The fur over the supraorbital rim was shaved and the mouse was placed on the stage of a soundproof enclosure. An MRI head holder was used to stabilize the position of the head. During the procedure, the mouse was continually supplied with vaporized isoflurane with oxygen. The microtip probe was placed against the supraorbital rim. The Sonifier was activated to deliver an 800 msec shock at a 35% or 40% amplitude (resulting in a 230–250-micron oscillation according to manufacturers’ specifications). The contralateral eye was used as a control. Following treatment, the mice were injected with bupronorphine for analgesia, placed in a new cage with thermal support, and monitored until they had fully recovered.

### Treatment with PEGylated arginase 1 (PEG-A1) and DFMO

For systemic treatment, PEG-A1 was administered by i.p. injection (25 mg/kg) at 3 h after injury and repeated every 3 days. PEG without A1 was used as vehicle control. Intravitreal delivery of PEG-A1 was performed following our established protocol [9]. PEG-A1 (1.7 ng/μL, 1 µl) or phosphate-buffered saline (PBS, 1 µl) was administered under anesthesia at 3 hrs. after injury via intravitreal injection using a Hamilton syringe. Some mice were also treated with DFMO (1 g/kg in PBS, i.p.) or vehicle (PBS) at 3 hr after ONC and supplied with 2% DFMO in their drinking water (or normal drinking water) until sacrifice.

### Visual function testing

Visual acuity and contrast sensitivity were assessed using optokinetic response tracking (OKT) (Cerebral Mechanics, Inc., Lethbridge, AB, Canada) as described previously [32]. RGC function was analyzed by pattern electroretinographic (PERG) recording according to an established protocol [33].

### Immunofluorescence imaging

Retinal flat mounts were prepared following our established protocol [34] and reacted with the following antibodies to detect RGCs, RGC axons, and microglia/MΦ [RNA-Binding Protein, mRNA Processing Factor (RBPMS), SMI-32, TUJI and anti-Iba1, respectively (**Table 1**)]. Next the samples were washed 3 times with PBS and reacted with goat anti-rabbit or anti-mouse secondary antibodies. Images were captured using a Zeiss Axioplan2 fluorescence microscope or Zeiss 780 inverted Confocal microscope (Carl Zeiss Meditec, Inc., Dublin, CA). For quantitation and analysis of RGC survival and axonal integrity, a series of nine images were taken of each retina using a 20X lens.

**Table 1.** Antibodies.

| **Antibody** | **Catalog number** | **Company** | **Dilution** | **Experiment** |
| --- | --- | --- | --- | --- |
| RBPMS | GTX118619 | GeneTex, Zeeland, MI | 1:200 | IF |
| Neurofilament H (SMI-32) | 801701 | Biolegend, San Diego, CA | 1:200 | IF |
| Tubulin $\beta$ 3 (TUBB3, TUJI) | 801202 | Biolegend, San Diego, CA | 1:200 | IF |
| Iba1 | 019-19741 | FUJIFILM Wako, Richmond, VA | 1:200 | IF |
| Glial Fibrillary Acidic Protein (GFAP) | Z0334 | Agilent Technologies, Santa Clara, CA | 1:200 | IF |
|  |  |  | 1:1000 | WB |
| Goat anti-Rabbit IgG (H+L), Alexa Fluor 488 | A-11034 | Thermo Fisher Scientific, Waltham, MA | 1:400 | IF |
| Goat anti-Rabbit IgG (H+L), Alexa Fluor 594 | A-11037 | Thermo Fisher Scientific, Waltham, MA | 1:400 | IF |
| Goat anti-Mouse IgG1, Alexa Fluor 568 | A-21124 | Thermo Fisher Scientific, Waltham, MA | 1:400 | IF |
| Goat anti-Mouse IgG2a, Alexa Fluor 594 | A-21135 | Thermo Fisher Scientific, Waltham, MA | 1:400 | IF |
| TNF-α | ab1793 | Abcam, Cambridge, United Kingdom | 1:500 | WB |
| TNFR1 | 3736 | Cell Signaling Technology, Danvers, MA | 1:1000 | WB |
| β-Actin | A1978 | MilliporeSigma, Burlington, MA | 1:2000 | WB |
| Rabbit IgG HRP | NA934 | Cytiva, Marlborough, MA | 1:2000 | WB |
| Mouse IgG HRP | NA931 | Cytiva, Marlborough, MA | 1:2000 | WB |

### Western blotting

Retinal proteins were extracted for Western blot analysis following our established protocol [35]. The samples were homogenized in modified RIPA buffer (20 mM Tris HCl, 2.5 mM EDTA, 50 mM NaF, 10 mMNa4P2O7, 1% Triton X-100, 0.1% sodium dodecyl sulfate, 1% sodium deoxycholate, 1 mM phenylmethylsulfonyl fluoride, pH 7.4). Samples containing equal amounts of protein were separated by 8%, 10% or 12% sodium dodecyl sulfate polyacrylamide gel electrophoresis, transferred to polyvinylidene difluoride (PVDF) or nitrocellulose membrane, and incubated overnight at 4°C with anti-GFAP, anti-TNF-α, and anti-TNFR1 as listed in **Table 1**. The following day, the membranes were washed 3 times with 1% TPBS and incubated with corresponding horseradish peroxidase-linked secondary antibodies (**Table 1**). Bands were quantified by densitometry and the data were analyzed using ImageJ software and normalized to loading controls [36]. Equal loading was verified by stripping the membranes and reacting them with a monoclonal antibody against β-actin.

### Quantitative RT PCR (qRT PCR)

Total RNA was extracted from retinal samples using an RNAqueous 4PCR total RNA isolation kit (Invitrogen, Carlsbad, CA, US) and qRT PCR was performed as described previously [19, 32]. Primer sequences for mouse transcripts are shown in **Table 2**. Levels of mRNA from sham, vehicle, and PEG-A1 treatment groups are shown as fold change relative to the controls

**Table 2.**
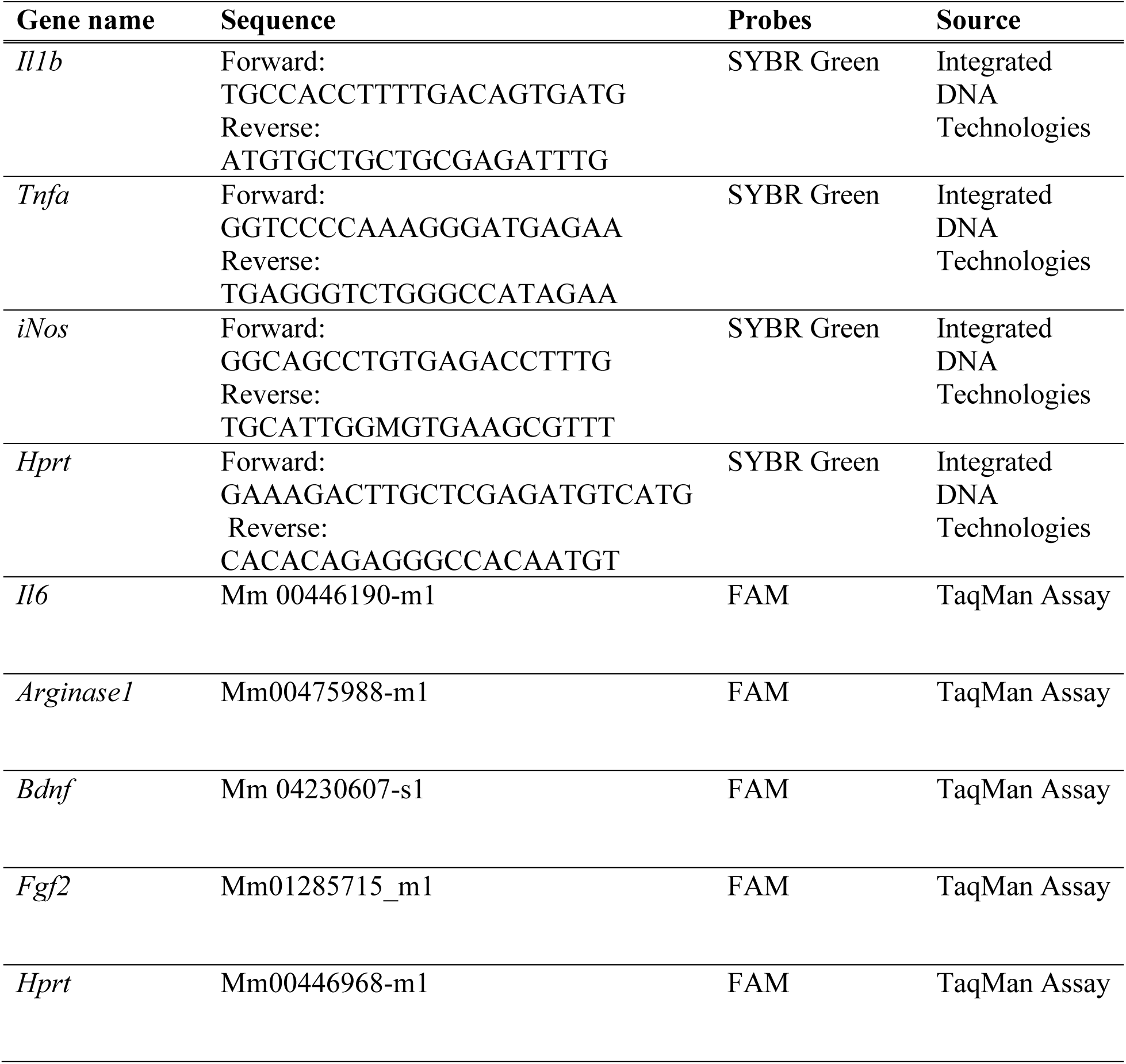
Primers.

### Liquid chromatography-multiple reaction monitoring mass spectrometry (LC-MRM MS)

Retinal samples were collected and the separation was performed using a Phenomenex Kinetex C18 column (100x2.1 mm, 1.7um) on a Shimadzu Nexera UHPLC system at a flow rate of 0.2 mL/min using gradient elution from 10% to 95% acetonitrile (with 0.1% formic acid) in 6 minutes. The effluent was ionized using positive ion electrospray on a TSQ Quantiva triple-quadrupole mass spectrometer with the following instrument settings: ion spray voltage 3500V, sheath gas 10, ion transfer tube temperature 350, aux gas 5, and Q1/Q3 resolution of 0.7 FWHM. The optimal collision energy and RF lens were determined using purchased standards. The integrated peak areas for the transitions were calculated for each sample using Skyline software (Version 20.0, University of Washington).

### Statistical analysis

Group differences were evaluated by using one-way analysis of variance of ranks. Results were considered significant at P<0.05. Data are presented as the mean ± SE.

## Results

### PEG-A1 treatment limits retinal injury in a mouse model of indirect TON

Our previous studies showed that arginase 1 (A1) deficiency worsens retinal inflammation and neurovascular injury in a mouse model of retinal I/R injury, demonstrating that A1 protects against retinal injury [9]. Furthermore, we found that systemic administration of PEG-A1 is neuroprotective in mouse models of retinal I/R injury and the ONC model of direct TON [19]. In the present study, we first evaluated whether PEG-A1 can also protect against neuronal injury and vision loss in the sonication-induced TON (SI-TON) model of indirect TON.

Mice underwent SI-TON, followed by intraperitoneal (i.p.) injection of PEG-A1 or control (PEG) at 3 hours after the injury, with subsequent administration every 3 days (**Fig. 1A**). Systemic administration via i.p. injection was used as we had previously demonstrated the protective effect of PEG-A1 in preventing retinal injury in models of ONC and ischemia reperfusion injury using this route of administration [9, 37]. Retinal flat-mount immunostaining was performed to quantify the numbers of RBPMS-positive RGCs in retinas collected at 7 days and 14 days after SI-TON. The vehicle-treated control group showed a significant loss of RGCs compared to the sham group, and this decrease in RGCs was significantly suppressed by PEG-A1 injection at both 7 and 14 days after the injury (**Fig. 1B, C, D, E**). These results indicate that systemic administration of PEG-A1 is neuroprotective after SI-TON.

**Fig. 1.**
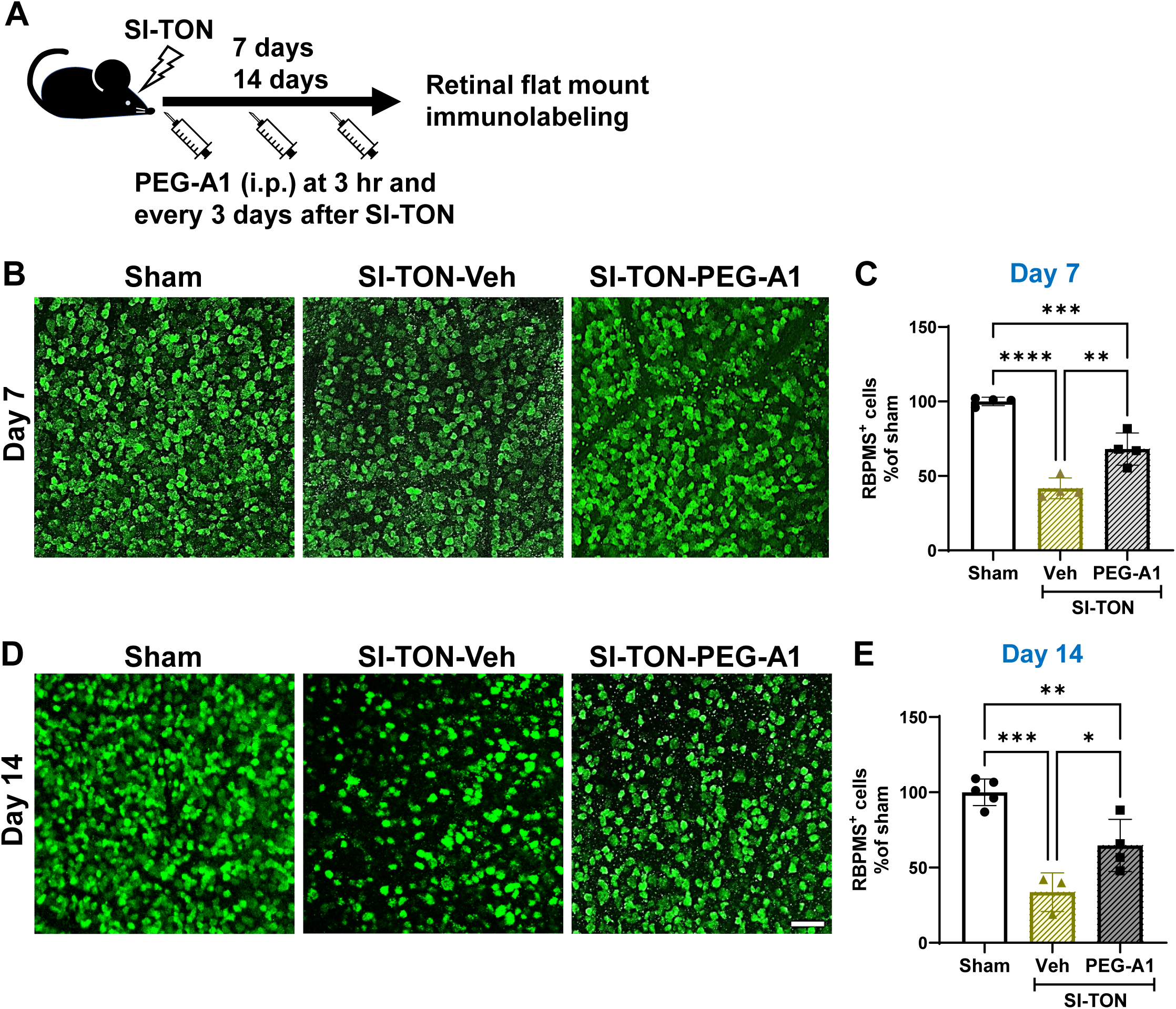
Intraperitoneal injection of PEG-A1 reduced SI-TON-induced neurodegeneration. (A) Schematic representation of the experimental protocol. Intraperitoneal injections of PEG-A1 were administered at 3 hr after SI-TON and repeated every 3 days. Retinal flat mounts were collected at day 7 and day 14 after SI-TON for immunostaining. Representative RBPMS labeling (B,D) and quantification (C,E) of retinal flat mounts show a significant decrease in RBPMS^+^ RGCs at 7 (B,C) and 14 days (D,E) after SI-TON. PEG-A1 treatment significantly reduced loss of RGCs as compared with vehicle. *N*=3-5. \**p*<0.05, \*\**p*<0.01, \*\*\**p*<0.001. Scale bar=50 μm

To assess the effect of local delivery of PEG-A1 on SI-TON-induced injury, we used intravitreal injections of PEG-A1 or PBS as vehicle control. A single intravitreal injection of PEG-A1 or PBS was administered 3 hours after SI-TON, and RBPMS immunostaining of retinal flat mounts was performed at 5 days after the injury (**Fig 2A**). The results showed that the number of RBPMS-positive RGCs was significantly reduced in the SI-TON vehicle control group compared to the sham group and that this cell loss was completely prevented in the SI-TON group treated with PEG-A1 (**Fig. 2B, C**). Since loss of the optic nerve axons is a major pathological event in TON, we also examined the effect of PEG-A1 on axonal survival after SI-TON. Quantification of SMI-32-positive RGC axons at 5 days after SI-TON using immunostaining of retinal flat mounts showed a significant reduction in numbers of surviving axons in the SI-TON vehicle control group compared to the sham group. This reduction was blocked by intravitreal administration of PEG-A1 (**Fig. 2D, E**). These results indicate that local administration of PEG-A1 prevents RGC neuronal and axonal damage caused by SI-TON, and that a single local administration preserves both RGCs and their axons.

**Fig. 2.**
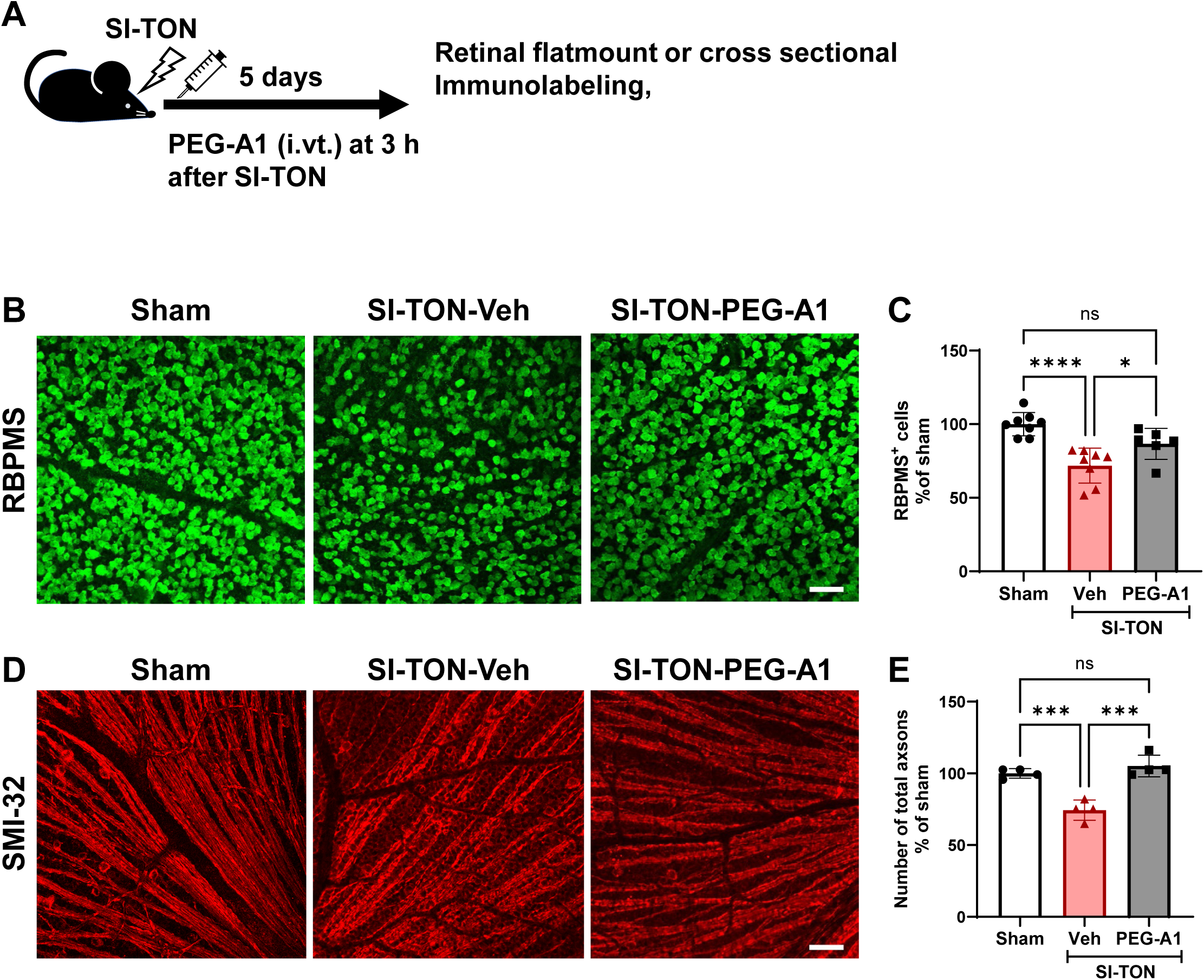
Intravitreal injection of PEG-A1 prevented SI-TON-induced neurodegeneration. (A) Schematic representation of the experimental protocol. Intravitreal injection of PEG-A1 or vehicle was performed at 3 hr after SI-TON. Experimental analyses were performed at 5 days after SI- TON. Representative RBPMS labeling (B) and quantification (C) of RBPMS^+^ cells in retinal flat mounts show a decrease in RGCs at 5 days after SI-TON. PEG-A1 treatment significantly preserved RGCs as compared with vehicle. *N*=6-8. \**p*<0.05, \*\*\*\**p*<0.0001, ns=not significant. Scale bar =50 μm. Representative SMI-32 labeling of RGC axons in retinal flat mounts (D) and quantification (E) show significant loss of RGC axons at 5 days after SI-TON. PEG-A1 treatment significantly preserved RGC axons as compared with vehicle. *N*=4. \**p*<0.05, \*\*\**p*<0.001, \*\*\*\**p*<0.0001, Scale bar=50 μm.

### PEG-A1 suppresses Műller glial and microglial cell activation and inflammatory cytokine production after SI-TON

We examined the effect of intravitreal injection PEG-A1 on inflammatory responses in Műller and microglial/MΦ cells after SI-TON. Immuno-labeling for GFAP, a marker of Műller cell activation, was performed on retinal cross sections. Results of this analysis showed high levels of GFAP immunoreactivity that extended from the ganglion cell layer (GCL) to the outer nuclear layer (ONL) in the SI-TON vehicle group, whereas GFAP immunoreactivity was restricted to the nerve fiber layer in the sham group (**Fig. 3A**). This increase in GFAP immunoreactivity was markedly suppressed by PEG-A1 treatment. Furthermore, Western blot analysis confirmed a significant increase in GFAP expression after SI-TON treatment and this increase was significantly inhibited by intravitreal administration of PEG-A1 (**Fig. 3B, C**). Immuno-labeling for Iba-1, a marker of microglia/MΦ, was performed to evaluate their activation after SI-TON. The SI-TON vehicle group showed a prominent increase in Iba-1-positive cells with an enlarged, activated morphology compared to the sham group, and this alteration was suppressed by PEG-A1 administration (**Fig. 3D, E**).

**Fig. 3.**
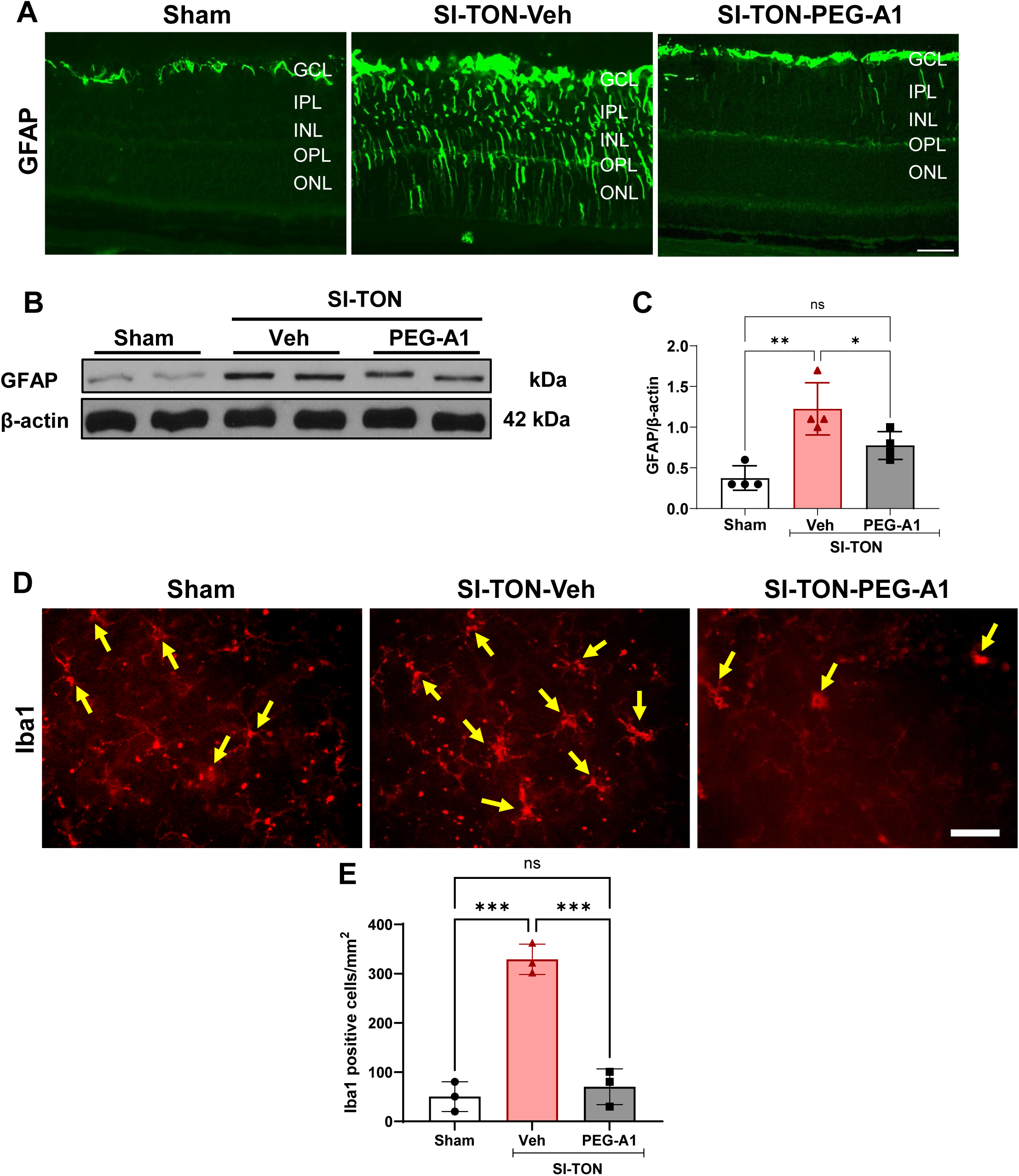
Intravitreal injection of PEG-A1 suppressed SI-TON-induced Müller glial and microglial cell activation. Representative GFAP labeling of retinal flatmounts **(A)**, Western blot analysis with retinal tissues **(B),** and quantification **(C)** show activation of Müller cells at 5 days after SI-TON. PEG-A1 treatment significantly inhibited Müller cell activation. *N*=4. \**p*<0.05, \*\**p*<0.01, ns=not significant. Scale bar=50mm. Iba1 labeling **(D)** of microglia/MΦ (arrows) and quantification **(E)** in retinal flatmounts show numerous activated profiles with enlarged cell bodies at 5 days after SI-TON. PEG-A1 treatment markedly reduced activated profiles as compared with vehicle. *N*=3. \*\*\**p*<0.001, ns=not significant. Scale bar=50 mm.

We also examined the effect of PEG-A1 treatment on SI-TON-induced inflammatory cytokine formation. Western blot analysis showed that SI-TON significantly increased the expression of tumor necrosis factor-α (TNF-α) and its receptor TNF-αR1. PEG-A1 treatment significantly suppressed these increases (**Fig. 4A-C**). Further analysis of mRNA levels by quantitative RT/PCR showed that SI-TON induced an increase in the expression of inflammatory markers, interleukin-1β (IL-1β), TNF-α, interleukin-6 (IL-6), and inducible nitric oxide synthase (iNOS) (**Fig. 4D**). These alterations were also significantly suppressed by PEG-A1 administration. In contrast, no significant difference in the expression of the growth/survival factors brain-derived neurotrophic factor (BDNF) and fibroblast growth factor 2 (FGF2) was observed between the sham and SI-TON groups, and no significant change was seen with PEG-A1 administration (**Fig. 4E**).

**Fig. 4.**
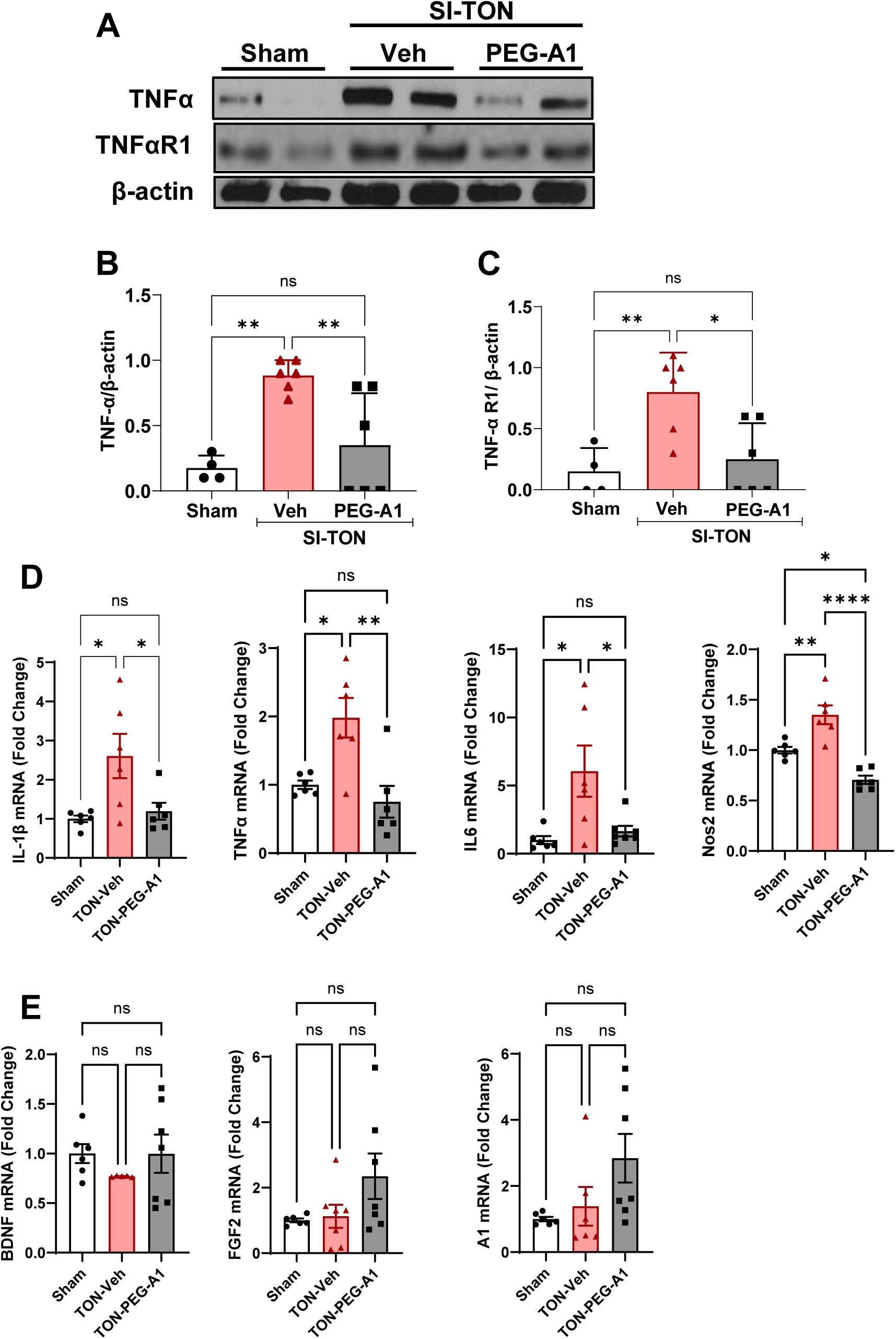
Intravitreal injection of PEG-A1 suppressed SI-TON-induced retinal inflammation. (A-C) Western blot analysis shows significant increases in TNF-α and TNFR1 in the vehicle group as compared with sham controls at 5 days after SI-TON. Intravitreal injection of PEG-A1 blocked these increases as compared with vehicle. *N*=4-6. \**p*<0.05, \*\**p*<0.01, ns=not significant. **(D)** qPCR shows significant increases in mRNA levels of proinflammatory markers at 5 days after SI-TON. These increases were blocked by PEG-A1 treatment. n=6. \**p*<0.05, \*\**p*<0.01, \*\*\*\**p*<0.0001, ns=not significant. **(E)** Neither SI-TON nor PEG-A1 altered mRNA levels of A1 or the anti-inflammatory markers BDNF and FGF2. *N*=6-7. ns=not significant.

### Intravitreal delivery of PEG-A1 preserves visual function following SI-TON

We examined the effect of PEG-A1 treatment on SI-TON-induced impairment of retinal and visual function. Analysis of the pattern ERG response, which reflects retinal ganglion cell (RGC) function, showed a significant reduction in the SI-TON group compared to the sham group (**Fig. 5A**). Although the PEG-A1-treated group showed a reduction in the Pattern ERG response compared to the sham group, this decrease was significantly less compared to the vehicle control group. Further analyses using the OptoMotry test showed that visual acuity and contrast sensitivity losses after SI-TON were also significantly improved by PEG-A1 administration as compared with the vehicle control group (**Fig. 5B, C**).

**Fig. 5.**
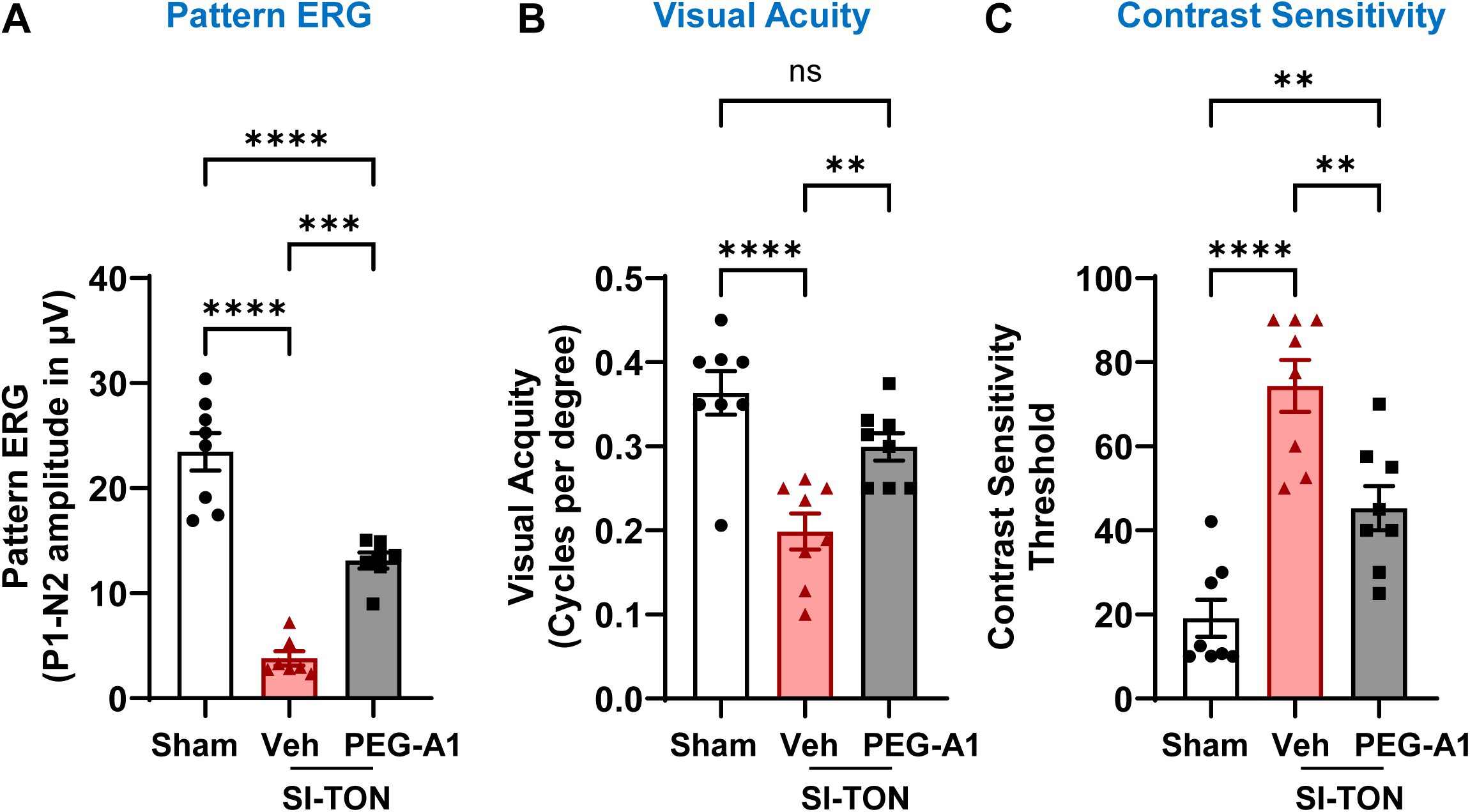
Intravitreal injection of PEG-A1 preserved retinal and visual function after SI-TON. Retinal function was measured by pattern ERG **(A)** and visual function was measured by OptoMotry responses **(B, C)** at 5 days after SI-TON injury. SI-TON injury decreased retinal and visual function, and both were significantly improved in the PEG-A1–treated group as compared with the vehicle group., *N*=7-8 **p<0.01, ****p<0.0001, ns=not significant.

### Intravitreal delivery of PEG-A1 limits neuronal injury following ONC

We evaluated the protective effect of intravitreal PEG-A1 delivery on retinal injury in the ONC model of direct TON. Retinal flat-mount immunostaining was performed to determine the treatment effects on RGCs and RGC axonal survival and microglia/MΦ activation in retinas collected at 7 days after ONC. ONC significantly reduced the number of RBPMS-positive RGCs (**Fig. 6B, C**) and TUJ1 positive axons (**Fig. 6D, E**) and significantly increased the number of Iba1-positive microglia/MΦ (**Fig. 6F, G**) as compared with the sham control and each of these changes was suppressed after intravitreal administration of PEG-A1. These results indicate that local delivery of PEG-A1 also protects against neuronal injury in the ONC model.

**Fig. 6.**
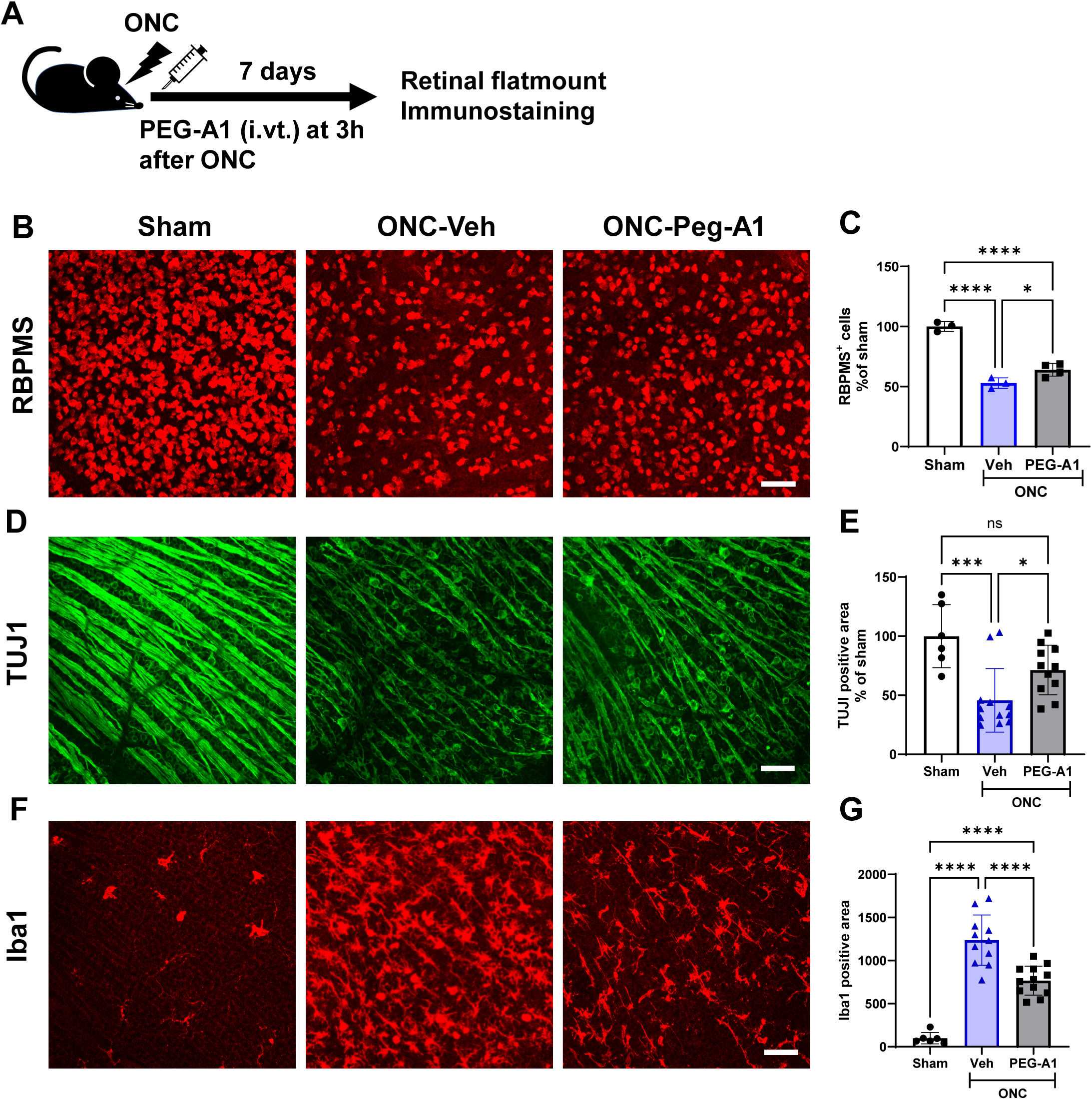
Intravitreal injection of PEG-A1 prevented ONC-induced neurodegeneration and microglial/MΦ activation. **(A)** Intravitreal injection of PEG-A1 or vehicle was administered at 3hr after ONC and retinas were collected at day 7 after ONC for immunostaining. Representative RBPMS labeling **(B)** and quantification **(C)** of RGCs in retinal flat mounts show a significant decline in RBPMS^+^ cells at 7 days after ONC. Single intravitreal injection of PEG-A1 significantly preserved RGCs as compared with vehicle. *N*=3-4. Scale bar=50 μm. Representative TUJ1 labeling **(D)** and quantification **(E)** of RGC axons in retinal flat mounts show a significant decrease in RGC axons after ONC. PEG-A1 treatment significantly preserved RGC axons as compared with vehicle controls. *N*=6-12. Scale bar=50 μm. Representative Iba1 labeling **(F)** and quantification **(G)** of microglia/MΦ in retinal flat mounts shows a significant increase in activated profiles after ONC. Intravitreal injection of PEG-A1 significantly reduced activated microglia/MΦ as compared with vehicle treatment. *N*=6-12. \**p*<0.05, \*\*\**p*<0.001, \*\*\*\**p*<0.0001. Scale bar=50 μm.

### The neuroprotective effect of PEG-A1 after ONC is blocked by inhibiting ODC

In order to determine whether the neuroprotective effect of PEG-A1 involved activation of the polyamine pathway, we treated the mice with DFMO, an ODC inhibitor, via intraperitoneal injection and in drinking water (**Fig. 7A**). First, we confirmed that DFMO alone had no effect on RGC survival. There was no difference in the number of RBPMS-positive cells between sham retinas with or without DFMO (**Fig. 7 B, C).** Similarly, DFMO did not affect RBPMS-positive cell number after ONC. Thus, DFMO alone does not affect RGC survival. Subsequently, we examined the combined effect of PEG-A1 and DFMO. As shown in Figure 7, the neuroprotective effects of PEG-A1 were suppressed by the DFMO treatment. These results indicate that activation of the polyamine pathway contributes to the neuroprotective effect of PEG-A1.

**Fig. 7.**
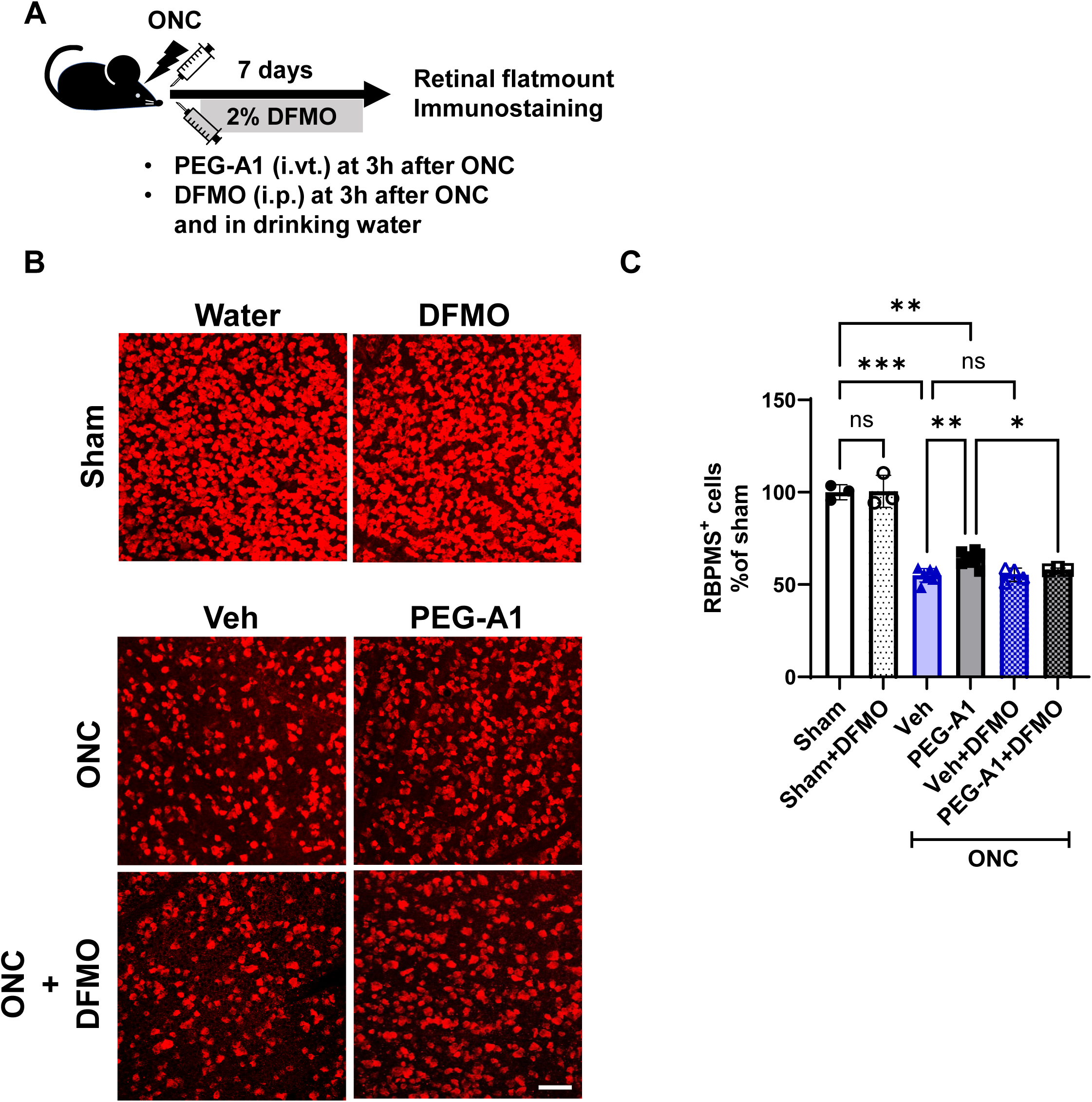
DFMO treatment abolished PEG-A1-mediated blockade of ONC-induced GCL loss. **(A)** Intravitreal injection of PEG-A1 or vehicle was performed at 3h after ONC and followed immediately by DFMO (1 g/kg, i.p.) or vehicle injection at 3h after ONC. Mice were supplied with 2% DFMO in drinking water or normal drinking water until day 7. Retinas were collected at day 7 after ONC for immunostaining. Representative RBPMS labeling **(B)** and quantification **(C)** of RGCs in retinal flat mounts show that there was no significant difference in number of RBPMS⁺ cells with or without DFMO treatment in sham retina. A marked decrease in RBPMS ^+^ cells was evident at 7 days after ONC. Intravitreal injection of PEG-A1 significantly improved RGC survival as compared with vehicle. DFMO treatment blocked the protective effect of PEG-A1. *N*=3-8. \**p*<0.05, \*\**p*<0.01, \*\*\**p*<0.001, ns=not significant. Scale bar=50 μm.

### PEG-A1 activates the polyamine pathway in the ONC model and this pathway is suppressed by DFMO

Because DFMO treatment blocked the neuroprotective effect of PEG-A1, we next analyzed changes in the levels of arginine/polyamine pathway metabolites including L-arginine, L-ornithine and polyamines (putrescine, spermidine and spermine) in the ONC model using LC-MS. There were no differences in their amounts between sham and ONC vehicle groups. (**Fig. 8A, B, C**). In contrast, PEG-A1 administration significantly increased L-ornithine and putrescine (**Fig. 8B, C**). These PEG-A1 induced increases were significantly suppressed by administration of DFMO. PEG-A1 treatment showed a trend towards increased spermidine and spermine levels, but the differences were not significant (**Fig. 8D, E)**.

**Fig. 8.**
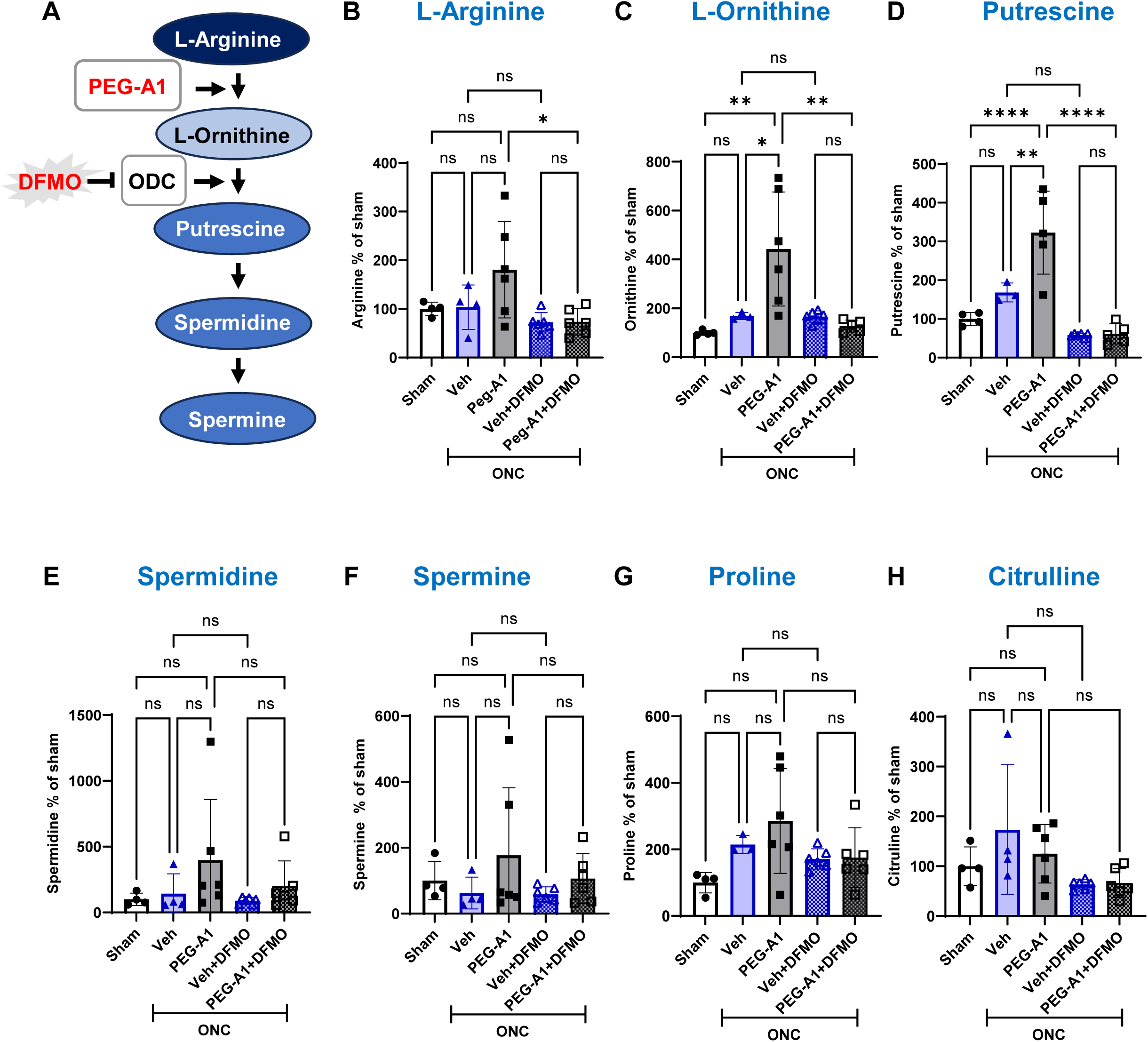
DFMO treatment abolished PEG-A1-mediated activation of the L-arginine/polyamine pathway. **(A)** Schematic illustration of the L-arginine-polyamine pathway regulated by PGC-A1 and inhibited by DFMO. **(B-H)** LC-MS analysis of retinal samples showed that intravitreal injection of PEG-A1 significantly increased L-ornithine and putrescine levels compared with sham and vehicle groups. DFMO or vehicle was administered at 3 hr after ONC, and mice were continuously supplied with 2% DFMO in drinking water or normal drinking water until sacrifice at day 3. PEG-A1 treatment significantly increased levels of L-ornithine and putrescine. DFMO treatment significantly reduced L-arginine, L-ornithine, and putrescine levels as compared with PEG-A1 treatment. *N*=4-6. \*\**p*<0.01, \*\*\*\**p*<0.0001.

Since L-ornithine is converted to proline via the ornithine aminotransferase pathway, we also evaluated the effects of PEG-A1 and DFMO on proline levels. Proline showed a tendency to increase in the ONC group compared to the sham group, but the difference was not significant (**Fig. 8F**). Furthermore, no significant effects of PEG-A1 or DFMO on proline levels were observed. We further examined changes in citrulline, an intermediate in NO synthesis pathway. Citrulline is generated during the conversion of substrate L-arginine to NO by NOS. Citrulline levels were not altered by ONC compared with sham and were not affected by PEG-A1 or DFMO (**Fig. 8G**). These results suggest that PEG-A1 activates the polyamine pathway, as evidenced by the increases in L-ornithine and putrescine, which were suppressed by administration of DFMO.

## Discussion

Optic neuropathy is associated with progressive secondary neurodegeneration and inflammatory responses after the initial injury. Effective neuroprotective therapies remain limited. In the present study, we investigated the neuroprotective effects of PEG-A1 in models of both indirect and direct optic neuropathy, the SI-TON and ONC models, respectively. Our studies focused on secondary injury-associated responses and L-arginine-related metabolic pathways. We found that PEG-A1 treatment after SI-TON improved survival of the retinal ganglion cells and their axons and also improved retinal and visual function while preventing activation of Muller glia and microglia/MΦ and limiting inflammation. PEG-A1 treatment also improved survival of retinal ganglion cells and their axons after ONC while reducing activation of microglial cells and MΦ. In addition, inhibition of ornithine decarboxylase activity blocked the PEG-A1-mediated neuroprotection, indicating functional involvement of this metabolic pathway in the neuroprotective effects. Furthermore, a single intravitreal administration of PEG-A1 was sufficient to confer neuroprotective effects in both SI-TON and ONC models.

PEG-A1 treatment inhibited SI-TON-induced injury as indicated by preservation of RBPMS-positive RGCs and conserved SMI-32-positive axonal integrity. Importantly, this structural preservation was accompanied by preservation of retinal and visual function, including improvements in the pattern ERG response, visual acuity, and contrast sensitivity as compared with the vehicle treated controls. These results suggest that PEG-A1 treatment attenuates SI-TON-associated neurodegeneration at both structural and functional levels. Because visual dysfunction after SI-TON is associated with progressive retinal ganglion cell and axonal loss [31, 38], the ability of PEG-A1 to preserve both neuronal structure and visual function suggests its potential as an effective therapy for traumatic optic neuropathy. However, the present study does not clarify whether PEG-A1 acts directly on neurons or preserves neuronal integrity indirectly through reduction of secondary injury-associated responses.

Our previous study using the ONC model showed that PEG-A1 treatment given by i.p. injection three hours after ONC crossed the blood retinal barrier, increased survival of NeuN-positive RGCs, limited the ONC-induced increases in Iba1-positive microglia/MΦ, and reduced IL-1β expression, while increasing IL-10 and BDNF expression [19]. Although ONC is an experimental model that directly induces optic nerve injury, secondary injury-associated inflammatory responses and glial activation also occur after the initial injury [5]. These previous results suggest that PEG-A1 treatment protects retinal neurons by limiting secondary injury-associated inflammatory reactions. Recent studies have shown that the arginase/ornithine decarboxylase pathway plays an important role in regulating MΦ inflammatory responses [20, 39]. Deletion of ornithine decarboxylase in macrophages has been shown to enhance their production of pro-inflammatory mediators, including IL-1β and TNF-α as well as increasing the expression of the iNOS gene Nos2 [26]. Consistent with these findings, our previous studies have shown that MΦ lacking arginase 1 have an increased inflammatory response to LPS stimulation *in vitro* [9]. Our present results show that PEG-A1 treatment reduced the activation of glia and microglial/MΦ as demonstrated by reductions in GFAP and Iba1 expression after SI-TON. PEG-A1 treatment also reduced TNF-α and TNFR1 protein expression and decreased levels of mRNA for Nos2 along with those for the inflammatory cytokines IL-1β, TNFα, and IL-6. In contrast, PEG-A1 treatment did not significantly change endogenous arginase 1, BDNF, or FGF2 expression in the SI-TON model. These findings suggest that PEG-A1 may mainly limit inflammatory signaling pathways without broadly altering injury-associated gene expression. However, our results do not clarify whether suppression of inflammatory and glial responses is a primary mechanism of neuroprotection or a secondary consequence of reduced neuronal injury.

PEG-A1 converts extracellular L-arginine into L-ornithine, thereby reducing the availability of L-arginine, as the substrate for iNOS, and thus inhibiting NO production. Previous studies in MΦ have shown that reduced L-arginine availability decreases iNOS expression and activity [40]. Our previous studies using MΦ from myeloid-specific arginase 1 knockout mice showed exaggerated inflammatory responses to LPS as indicated by increases in their expression of iNOS and other inflammatory mediators, whereas PEG-A1 treatment suppressed those effects [9]. In the present study, PEG-A1 treatment reduced Nos2 mRNA expression. This suggests that in addition to the suppression of iNOS activity due to a decrease in L-arginine, a reduction in the inflammatory response in MΦ via ornithine/polyamine metabolism or a decrease in microglia/MΦ activation may have contributed to the drop in Nos2 expression. Therefore, PEG-A1 may contribute to neuroprotection by reducing nitrosative/oxidative stress mediated by the iNOS/NO pathway.

The present study further provides mechanistic evidence that activation of the arginase-ornithine-putrescine pathway contributes to PEG-A1-mediated neuroprotection. PEG-A1 treatment after ONC also changed metabolites related to L-arginine and polyamine metabolism, significantly increasing L-ornithine, a direct product of arginase activity, as well as putrescine, a downstream L-ornithine metabolite. Previous studies have shown that putrescine suppresses inflammatory gene expression in activated microglia/MΦ [26, 41]. Consistent with that report, PEG-A1 treatment reduced the number of Iba1 positive cells in both ONC and SI-TON models, supporting the idea that activation of the arginase-ornithine-putrescine pathway is associated with suppression of inflammatory cell activation. Furthermore, inhibition of ornithine decarboxylase with DFMO blocked the increase in putrescine and limited the neuroprotective effect of PEG-A1. These findings suggest that activation of the ODC-dependent arginase-ornithine-putrescine pathway is required for PEG-A1 mediated neuroprotection.

Interestingly, the role of polyamine metabolism in neuronal injury appears to be highly context dependent. Previous studies have reported that pretreatment with DFMO can protect retinal neurons in models of NMDA-induced excitotoxicity [42]. Pretreatment with a polyamine oxidase inhibitor has also been shown to prevent excitotoxicity-induced retinal neurodegeneration [43] These results suggest that polyamines may contribute to NMDA receptor-mediated neuronal injury under some conditions. In contrast, our present results showed that inhibiting the L-ornithine-polyamine pathway with DFMO abolished the neuroprotective effect of PEG-A1. While polyamines have been implicated in excitotoxic signaling and the generation of toxic oxidative metabolites through polyamine catabolism, they have also been associated with tissue repair and neuroprotection [44]. Therefore, the effects of DFMO on excitotoxic neuronal injury may depend on the timing of treatment and the stage of injury. Further studies are needed to determine how polyamine metabolism influences acute excitotoxic injury versus later secondary injury responses following optic nerve trauma.

The present study also demonstrated that a single intravitreal administration of PEG-A1 was sufficient to confer neuroprotective effects after either SI-TON or ONC injury. Intravitreal PEG-A1 treatment preserved RGCs and axonal integrity, limited microglial/MΦ activation, and improved visual function after SI-TON. In addition, intravitreal PEG-A1 treatment also preserved RGCs and their axons and reduced microglial/MΦ activation in the ONC model. Clinical studies have shown that repeated systemic administration of PEG-A1 is generally well tolerated, although treatment-related adverse events have been reported [12, 14]. In the present study, neuroprotective effects were achieved with a single intravitreal administration of PEG-A1, suggesting that local ocular delivery could reduce the risk of systemic adverse events while maintaining therapeutic efficacy.

In summary, the present study demonstrated that PEG-A1 exerts neuroprotective effects after SI-TON and ONC and suppresses injury-associated inflammation and glial and microglial/MΦ activation. PEG-A1 treatment also altered metabolites associated with the arginine-ornithine-putrescine axis, and inhibition of the ornithine/polyamine pathway blocked PEG-A1-mediated neuroprotection, suggesting involvement of polyamine-related metabolic pathways in the protective effects of PEG-A1. In addition, the present findings showed that a single intravitreal injection of PEG-A1 was sufficient to confer neuroprotection after either SI-TON or ONC. Together, these findings suggest that PEG-A1 treatment could offer a therapeutic strategy for limiting retinal injury during TON through modulation of secondary injury-associated responses and L-arginine-related metabolic pathways.

## Abbreviations

A1: Arginase 1
A2: Arginase 2
BDNF: Brain-derived neurotrophic factor
DFMO: Difluoromethylornithine
ERG: Electroretinogram
FGF2: Fibroblast growth factor 2
GFAP: Glial fibrillary acidic protein
IL: Interleukin
I/R: Ischemia/reperfusion
MΦ: Macrophages
NMDA: N-methyl-D-aspartate
NOS: Nitric oxide synthase
ODC: Ornithine decarboxylase
ONC: Optic nerve crush
PEG: Polyethylene glycol
RBPMS: RNA-binding protein with multiple splicing
RGC: Retinal ganglion cells
SI-TON: Sonication-induced TON
TNF-α: Tumor necrosis factor-α
TON: Traumatic optic neuropathy
WT: Wild type

## Ethics declarations Ethics approval and consent to participate

All animal procedures complied with the Public Health Service Policy on Humane Care and Use of Laboratory Animals (revised July 2017) and with the statement from the Association for Research in Vision and Ophthalmology (ARVO) for the use of animals in ophthalmic and vision research. Procedures were approved by the institutional animal care and use committee (Animal Welfare Assurance # D16-00197).

## Consent for publication

Not applicable.

## Availability of data and materials

The data that support the findings of this study are available from the corresponding authors upon reasonable request.

## Authors’ contributions

M.Y., S.A.H.Z., M.A.R., R.W.C., and R.B.C. designed the study; T.L., Z.X., M.Y., S.A.H.Z., P.V.S., and M.A.R. performed/assisted research; S.A.H.Z., and M.A.R. analyzed the data; M.Y., S.A.H.Z., M.A.R., R.W.C., and R.B.C. provided inputs on experimental design and data interpretation and edited the manuscript; M.Y. and M.A.R. wrote the manuscript. All of the authors reviewed the manuscript.

## Acknowledgements

Not applicable

